# Exploring the Potential of AlphaFold Distograms for Predicting Binding-induced Hinge Motions

**DOI:** 10.1101/2025.07.25.666757

**Authors:** Büşra Savaş, Ayşe Berçin Barlas, Ezgi Karaca

**Affiliations:** İzmir Biomedicine and Genome Center, İzmir, Türkiye; Izmir International Biomedicine and Genome Institute, Dokuz Eylül University, Izmir, Türkiye

**Keywords:** AlphaFold, Distogram, Conformational flexibility, Binding-induced hinge motion, Multiple sequence alignment

## Abstract

AlphaFold models provide static structural predictions, limiting their use in interpreting flexible regions in low-resolution cryo-EM maps. Here, we assess whether AlphaFold-generated distograms can instead reveal conformational flexibility, focusing on binding-induced hinge motions. For this, we examined the key metabolic AK2/AIFM1 complex, where molecular dynamics and cryo-EM confirm a hinge motion in AK2 upon binding. Notably, this motion is captured in the AlphaFold2/3 distograms of apo AK2, even though it is absent in the predicted structures. By extending our analysis to other systems, we demonstrate that distograms may offer a valuable, model-independent method for interpreting ambiguous regions in cryo-EM maps.

## Introduction

Before the AlphaFold2 (AF2) revolution, modeling protein interactions within low-resolution cryo-electron microscopy (cryo-EM) maps predominantly relied on integrative structural biology approaches [1–5]. The use of AF2 significantly advanced this process by enabling the fitting of accurate structure predictions into cryo-EM maps [6–8]. Despite these advances, AF2’s utility remains limited when it comes to interpreting ambiguous regions in cryo-EM densities, caused by conformational flexibility [7, 9–12]. This is particularly pronounced in metabolic complexes, where functional transitions often involve hinge motions upon partner binding [6, 13–21]. To address this drawback, several recent methods have sought to repurpose AF2 as a conformational sampler [22–26]. One class of such methods manipulates the input multiple sequence alignment (MSA) while masking specific residue positions (columns) or entire sequences (rows) to encourage alternative conformational sampling [27,28]. Another strategy uses template-based guidance, where homologous structures from distinct functional states are used to steer predictions toward open or closed conformations [29,30]. Though, both strategies face practical limitations: MSA manipulation can lead to sampling of non-physiological conformations and appropriate templates are often unavailable. Moreover, these methods lack an intrinsic mechanism to evaluate which conformation best represents the biologically relevant state.

Before AF2’s widespread use, another source to explore protein flexibility was proposed by Schwarz *et al.*. In 2021, they demonstrated that conformational variability at the residue level could be inferred from the shape of inter-residue distance probability distributions, known as distograms [31]. Introduced in CASP13 by AlphaFold1, distograms provide probabilistic distance profiles between all residue pairs [32]. Using DMPFold to generate distograms, Schwarz *et al.* demonstrated that residue pairs with known conformational variability exhibited multimodal distance distributions, while rigid regions showed unimodal sharp peaks [31,33]. These findings suggest that the number and shape of peaks in a distogram may serve as practical indicators of conformational flexibility, while utilizing the full MSA information. Building on this insight, we investigated whether distograms, generated by AF2 and AF3 can hint at binding-induced hinge motions, an integral change in metabolic complexes.

To explore this hypothesis, we selected a recent cryo-EM structure of a complex formed between adenylate kinase 2 (AK2) and apoptosis-inducing factor mitochondria-associated 1 (AIFM1) as a test case [34]. This key mitochondrial assembly is involved in apoptotic signaling and ATP homeostasis. This system serves as an ideal proof-of-concept, as AK2 undergoes a distinct binding-induced hinge motion not observed in other adenylate kinases. Here, we combined AF modeling with molecular dynamics (MD) simulations to compare predicted and observed inter-residue distance probability distributions, focusing on the hinge motion [7,35]. To evaluate the broader applicability of this approach, we extended our analysis to three additional protein systems that may undergo binding-induced hinge motions.

Collectively, our findings provide preliminary yet promising evidence that AF-generated distogram profiles can serve as effective, structure-free indicators of hinge-based motions, potentially aiding in the interpretation of ambiguous regions in cryo-EM maps of metabolic complexes.

## Results

### A Key Metabolic Complex as a Proof-of-Concept for Distogram-Based Hinge Motion Prediction

AK2 is a mitochondrial isoform of the adenylate kinase enzyme family, responsible for maintaining cellular energy homeostasis. It catalyzes the reversible phosphorylation of AMP to ADP by using ATP [36,37]. As such, AK2 controls adenine nucleotide levels in the intermembrane space. By interacting with AIFM1, AK2 modulates redox-sensitive conformation of AIFM1, acting as a molecular switch between metabolic homeostasis and caspase-independent cell death [36–40]. Beyond its well-characterized domain motions, AK2 also exhibits a binding-induced hinge motion that is not captured in the previously resolved structures of this family. Two recent cryo-EM structures of the human AK2/AIFM1 complex reveal that the flexible C-terminal region of AK2, which is unresolved in the crystal structure of apo AK2 (PDB ID: 2C9Y), adopts a β-strand stacking against the C-terminal domain of AIFM1 [34,41,42]. In one of these structures, Schildhauer *et al.* used AF3 to model the complex and successfully fit the predicted β-stacking into their EM density (PDB ID: 9FL7; overall resolution: 4.3 Å; local resolution at the stacking site: ∼5 Å) [34]. This finding was independently validated by Rothemann *et al.*, where the last eight C-terminal residues of AK2 are resolved, forming the β-stacking interface (PDB ID: 9GR0; overall resolution: 2.6 Å; local resolution: up to 3.2 Å) [41].

The unique C-terminal rearrangement observed in AK2 presents an unexplored opportunity to evaluate whether this specific binding-induced hinge motion can be captured by different AF versions. Expanding on this, we researched initially how AF3 handles the flexible C-terminal region of AK2 and observed that AF3 predicted it as an α-helix with low confidence, reflected by a mean pLDDT score of 54 (Figure 1A) Furthermore, AK2’s PAE (predicted alignment error) matrix revealed that this C-terminal segment behaves as an independently moving domain (Figure S1). As it was already reported, AF3 modeled the C-terminal β-strand stacking against AIFM1 with high confidence (mean pLDDT: 77.6; Figure 1B) [34]. A comparison of the predicted apo and holo models showed a substantial conformational shift at the C-terminus, with a root mean square deviation (RMSD) of 12 Å across the terminal residues. In particular, the Cβ–Cβ distance between residues 230 and 233 increased from 6.2 Å in the apo model to 10.5 Å in the holo model, capturing a key hinge motion. Because this distance best reflects our focus on binding-induced conformational change (Figure S2), we use the 230–233 residue pair as a marker to track this hinge motion throughout the study.

**Figure 1.**
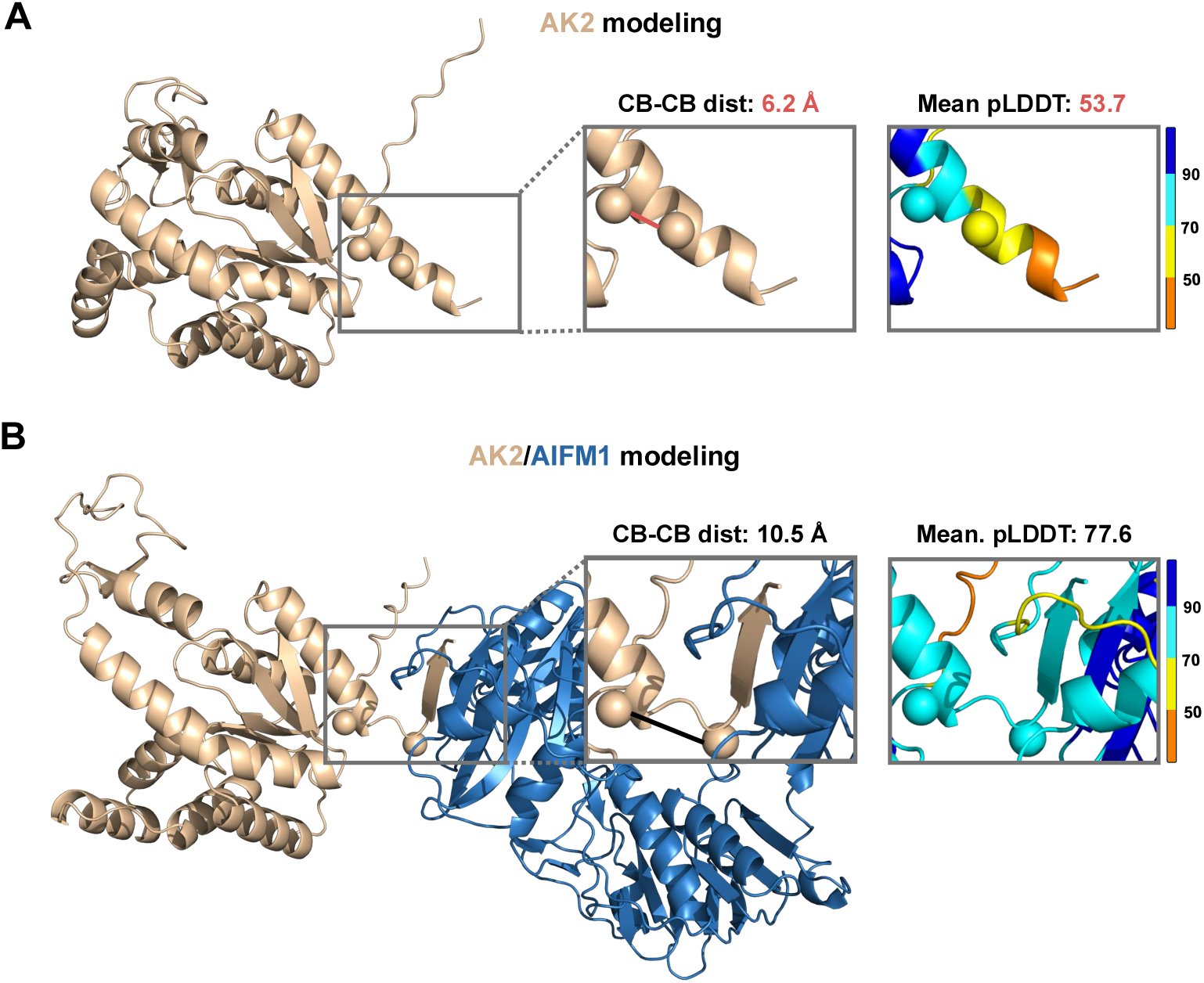
AlphaFold3 models capture a hinge-driven conformational switch of AK2. Top-ranked AF3 predictions (among 25 models) for apo **(A)** and holo **(B)** states of AK2 are shown. AK2 is depicted in wheat and AIFM1 in blue. Residues 230 and 233 of AK2, located at the flexible C-terminal end, are highlighted as spheres. Insets show close-up views of the Cβ–Cβ marker distance, together with the mean Cβ pLDDT scores of the last ten residues. pLDDT coloring indicates model confidence, with blue representing high and orange low confidence.

As a reference for ground truth, we monitored this marker distance between residues 230 and 233 with MD simulations. For the holo state, we relied on the previously published 4 μs all-atom MD trajectories of the AK2/AIFM1 complex (Figure S3) [34,43]. To provide a comparative baseline, we conducted 4 μs of MD simulations for the apo AK2. In the apo simulations, the marker distance predominantly ranged between 5.0 Å and 10.2 Å, with only ∼1.7% of frames exceeding 7.6 Å, which is the minimum distance observed in the holo trajectory (Figure 2A left, Figure S3). In contrast, the holo simulations showed a broader distribution from 7.6 to 13.3 Å, with a clear peak near 10 Å, consistent with the stabilized β-stacking conformation reported earlier (Figure S3). For subsequent analysis, we used the median Cβ–Cβ distances from the MD simulations as reference thresholds: 6.0 Å (with a probability of 0.4) for the apo state and 10.0 Å (with a probability of 0.2) for the holo state.

**Figure 2.**
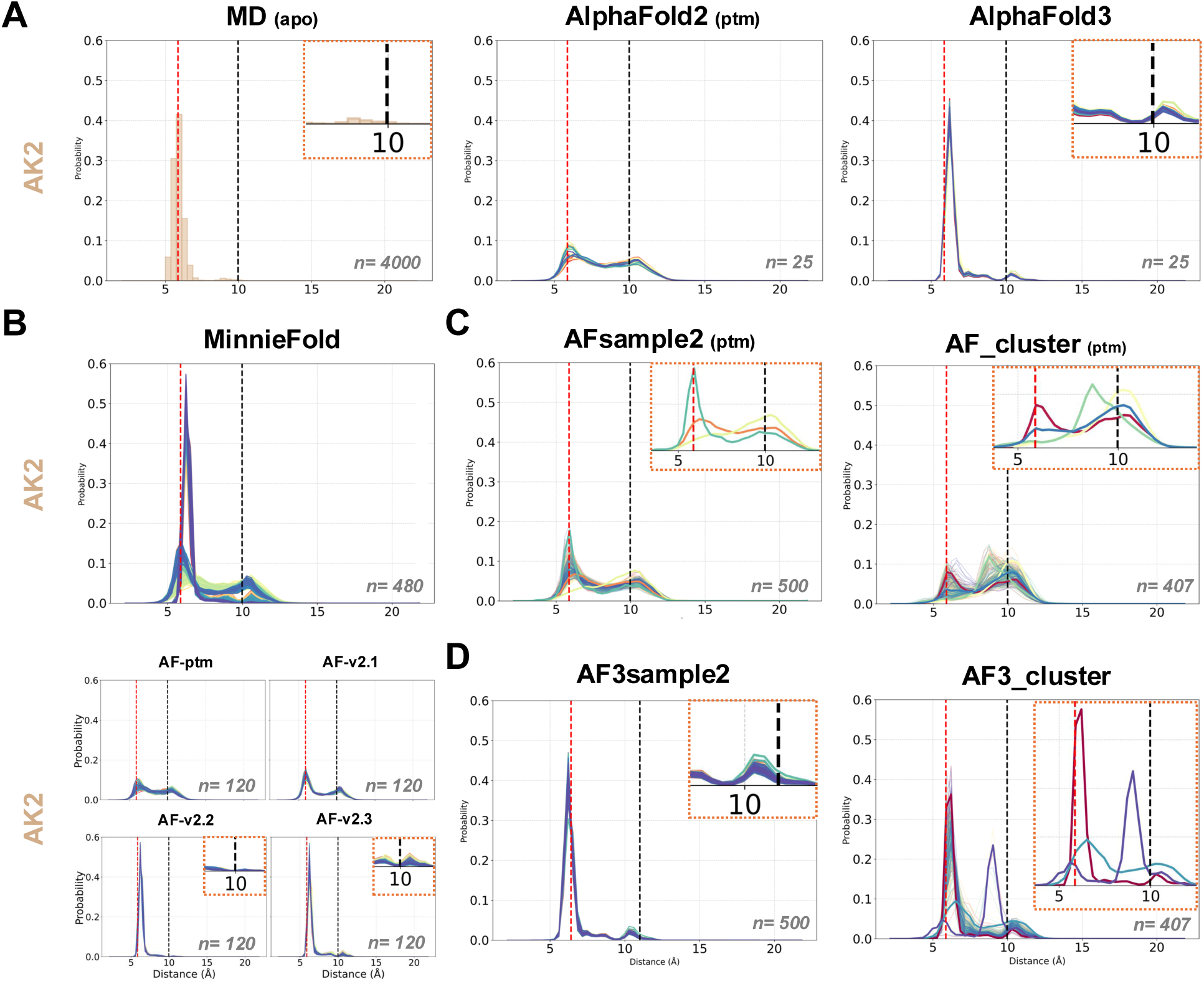
Comparison of marker hinge motion distances for apo AK2. Distogram profiles of the predicted 230-233 marker distance of apo AK2. Left: The measured distance probability distribution, obtained from apo MD simulations. Middle: AF2 distogram for the same pair. Right: AF3 distogram for the same pair. Distogram profiles of 230-233 pair calculated by (**B)** MinnieFold; all (top) and by version (bottom), (**C)** AFsample2 and AF_Cluster, (**D**) AF3sample2 and AF3_cluster. The dashed lines indicate the threshold distances obtained from MD simulations: at 6.0 Å in red for the apo and 10.0 Å in black for the holo state. Close-up views are highlighted with orange dashed frames. The sample given each plot is provided on the panel, indicated with *n*.

### Distogram Profile Evaluation Across Different AlphaFold Versions

To assess how accurately different AF versions reflect the binding-induced hinge dynamics captured in our MD simulations, we compared our MD-based marker distance distributions with those predicted by the distograms of AF2 and AF3. For this, we generated 25 independent distance probability distributions by each AI model without the use of template with the identical MSA inputs. The MSA was constructed with MMseqs2 through ColabFold, where the model generation followed the default settings of AF2.0 monomer (alphafold_ptm) and the latest version of AlphaFold, i.e., AF3 (see Supplementary Text) [44,45]. As an outcome, in the apo state, AF2-ptm generated broad, bimodal distograms with peaks centered near ∼6 Å and ∼10 Å (Figure 2A, middle). These peaks correspond approximately to the median distances observed in apo and holo MD simulations, respectively. However, the distinction between the two peaks was limited, with the holo-associated peak reaching a probability of no more than 0.1. In contrast, AF3 produced distograms that closely mirrored the apo MD distribution in both peak positions and relative amplitudes, showing a striking agreement with the simulation data (Figure 2A, right).

To further examine how different AF2 enhanced sampling configurations influence distogram outputs, we used our in-house sampling pipeline, MinnieFold, which systematically explores structural variability across four major AF2 versions (ptm, v2.1, v2.2, and v2.3) [46–48]. Inspired by the Massive Sampling approach [25], MinnieFold systematically varies the key AF2 sampling parameters such as the number of recycling steps, dropout settings, and template inclusion. It generates a total of 480 models across all four AF2 versions, enabling broad sampling of structural diversity in a computationally efficient manner (Supplementary Text). In the blind protein complex prediction challenge CAPRI55, we demonstrated that MinnieFold can produce interaction ensembles up to 11.7 Å RMSD variation. In this work, we applied MinnieFold to investigate how enhanced sampling affects distogram outputs, using the same input MSA as in our previous runs. The resulting distance distributions showed a mix of peak profiles, combining features seen in both AF2-ptm and AF3 (Figure 2B-top). When analyzed individually (Figure 2B-bottom), we saw that ptm and v2.1 displayed similar broad bimodal profiles, with v2.1 showing a stronger apo-associated peak, indicating a tendency toward the unbound state. In contrast, v2.2 produced sharply unimodal distributions centered around 6.0 Å, with a high probability (∼0.6). Notably, v2.3 reproduced a bimodal distribution closely matching the MD and AF3 profiles. These results suggest that increased sampling through MinnieFold does not introduce new flexibility signals beyond those observed in standard AF2 runs, but it can help amplify existing conformational features already encoded in specific versions.

### Distogram Profile Evaluation by Using Different MSA Content

Since parameter tuning alone did not introduce substantial diversity in the distogram outputs of AF2, we examined the influence of the input MSA on the breadth of distograms. For this, we first used AFsample2, which introduces stochasticity upon randomly masking MSA columns to perturb coevolutionary signals [27]. The second approach was AF_cluster, which clusters sequences to build reduced MSAs that represent distinct evolutionary subspaces [28]. Both strategies were implemented using the AF2-ptm model for consistency. In the case of AFsample2, the overall bimodal character of the distograms was preserved, but the probability of the apo-associated distance peak approximately doubled compared to the original AF2 run (Figure 2C). In contrast, AF_cluster produced a distinct unimodal peak centered around 8 Å with a probability of approximately 0.15, in addition to the original apo and holo peaks observed in AF2 (Figure 2C, right). This intermediate peak may represent a false positive, resulting from reduced MSA depth or limited sequence diversity, or it could reflect a true transition state between the apo and holo conformations. We performed the same protocols with AF3 with the same MSA inputs (Figure 2D, see Supplementary Text). Similar to AF_cluster, clustered MSA in AF3 produced distinct variations in peak positions, whereas column masking did not substantially affect the peak profiles.

### The Influence of MSA Pairing and Iterative Refinement on the Distogram Profiles

Since we unexpectedly observed that AF distograms successfully capture binding-induced hinge motion of AK2 from its apo state, we wanted to further investigate how evolutionary information derived from its binding partner would influence AK2’s distogram profiles. For the complex prediction, MSAs can be constructed either by pairing sequences of interacting proteins to preserve potential coevolutionary signals or by generating separate alignments for each protein partner [49]. To assess how this choice impacts the modeling process, we compared the distogram profiles of different AF2 versions using paired and unpaired MSAs for the modeling of AK2/AIFM1 complex (Figure S4).

When AK2/AIFM1 evolutionary information was provided as paired sequences, AF2 multimer (v2.1-3) marker distance distograms retained a bimodal character. In contrast, when unpaired MSAs were used, AF2.2 and AF2.3 distograms converged to a unimodal distribution centered around the holo-state median (∼10Å) (Figures S2, S4). A recent work on protein-peptide modeling presented a similar observation, where Guan and Keating showed that AF2 multimer versions can find the correct interchain coevolutionary signals from unpaired MSAs [50]. Interestingly, AF2-ptm showed the opposite trend, with unpaired MSAs producing a bimodal distribution and paired MSAs reducing this bimodality toward the holo-state distance. These pairing effects likely arise because AF2-ptm is trained on single-chain inputs, whereas AF2.2-3 is trained for complexes.

To gain further insight into what drives the transition from bimodal to unimodal distributions, we analyzed how the distograms evolved across recycling iterations. Each recycle updates pair representations based on inter-residue distance predictions, refining the model incrementally. In AK2/AIFM1 complex (Figure S5), the probability density around the holo-associated peak gradually increased with recycling, while the apo-like peak diminished (when AF2-ptm is used). This progressive shift suggests that the model incrementally refines its internal representation when the binding partner is present, leading to a more stable conformation seen in the holo state.

### Translation of Distogram Probabilities into Structural Ensembles

While both apo and holo distograms successfully captured the conformational landscape observed in MD simulations for the marker distance, a key question remained: to what extent are these probability distributions are reflected in the final predicted structures? To answer this, we analyzed the marker Cβ–Cβ distance in the structural ensembles generated by each of the previously described methods (Figure 3). Consequently, most of the methods, specifically AF2, MinnieFold, AFsample2, AF3, and AF3sample2 produced predicted structures that exhibit a sharply unimodal distance distribution concentrated tightly around the 6.0 Å apo distance. In contrast, the structural ensembles produced by the MSA clustering strategies did not exhibit this unimodal convergence. AF_cluster outputs spanned a broad, bimodal distribution resembling that of the AF2-ptm distogram, whereas AF3_cluster exhibited a dominant apo-state distance and a minor holo-state population, consistent with its underlying AF3 distogram. We also generated conformational ensembles using BioEmu, Boltz-2, and Chai-1 (see Supplementary Text) to evaluate whether their generative frameworks could more effectively capture conformational diversity [51–53]. All three methods produced significantly broader, bimodal distance distributions, successfully sampling both the apo and holo states. Notably, BioEmu, which integrates AF2-ptm pair embeddings within a diffusion-based framework, yielded ensemble distributions closely matching the AF2-ptm distogram. Interestingly, Boltz-2 produced the ensemble most consistent with the MD-derived distance distribution. Specifically, Boltz-2 uses ensemble-based supervision by integrating multiple samples from experimental ensembles and MD trajectories, suggesting it as a promising alternative in conformational sampling.

**Figure 3.**
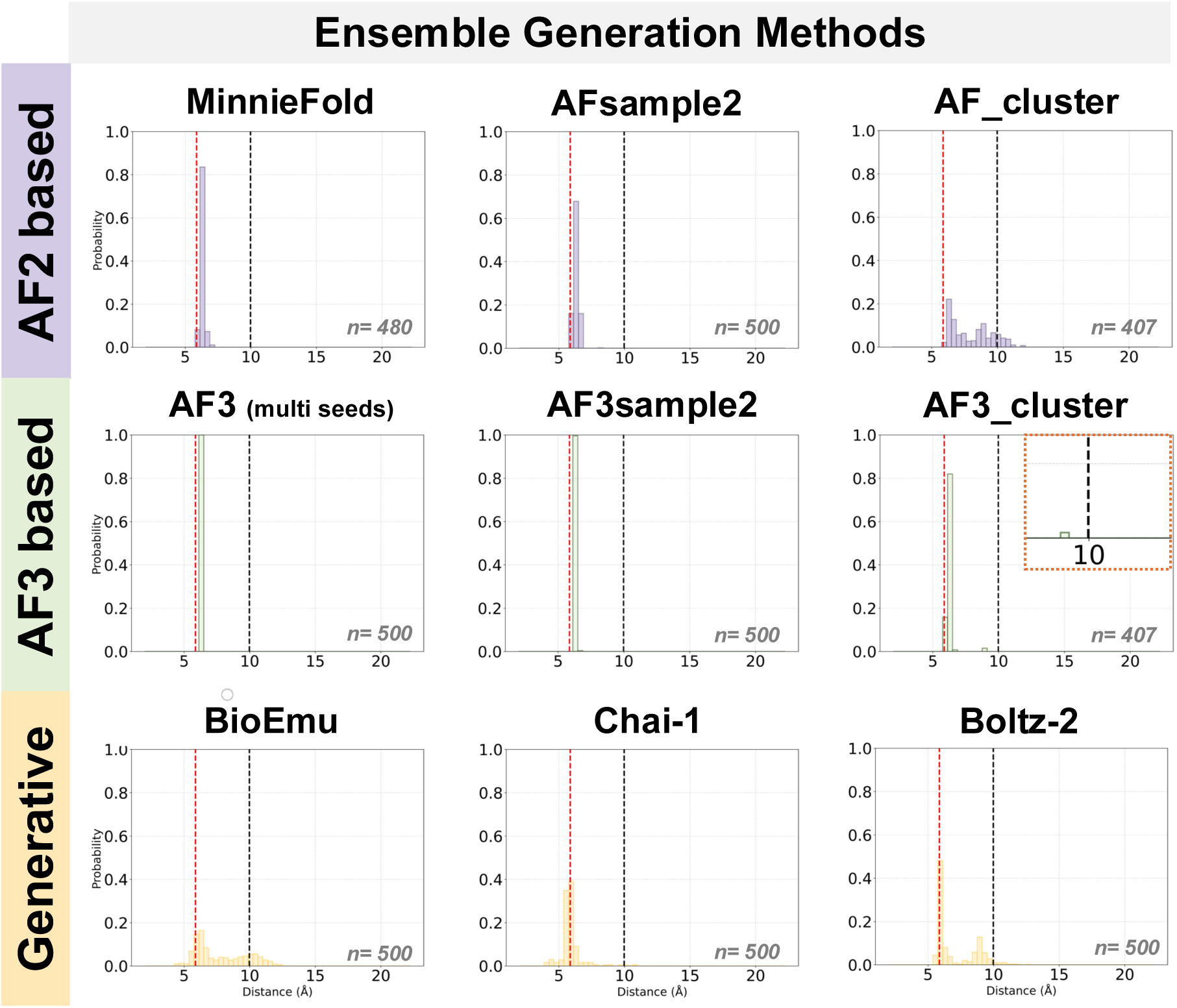
Comparison of marker distance profiles of AlphaFold2, AlphaFold3, and generative model-based sampling methods in capturing conformational diversity of AK2. The dashed lines indicate the threshold distances obtained from MD simulations: at 6.0 Å in red for the apo and 10.0 Å in black for the holo state. The sample given each plot is provided on the panel, indicated with n.

### Evaluating the Generalizability of Distogram Flexibility Signals

To test the generalizability of our distogram findings beyond the AK2/AIFM1 system, we analyzed three proteins with varying structural contexts: IL1R1, MIA40, and EIF2B3 (Figure 4). IL1R1 exhibits known binding-induced hinge motion, while MIA40 is predicted to undergo N-terminal relocation. Additionally, the EM map of EIF2B3 reveals an unresolved C-terminal peptide potentially indicative of hinge motion. For all the systems, we generated all-to-all residue distograms to identify bimodal profiles in regions of interest (Figure S6A-C), defined marker residue pairs, and analyzed distogram profiles across all standard AF versions (Figure 4). Consistent with our AK2 analysis, we also employed MinnieFold, MSA perturbation, and generative models to assess their capacity in predicting the expected conformational changes (Figure S7-9). Building on these benchmarks, the following sections evaluate the generalizability of our approach across these systems.

**Figure 4.**
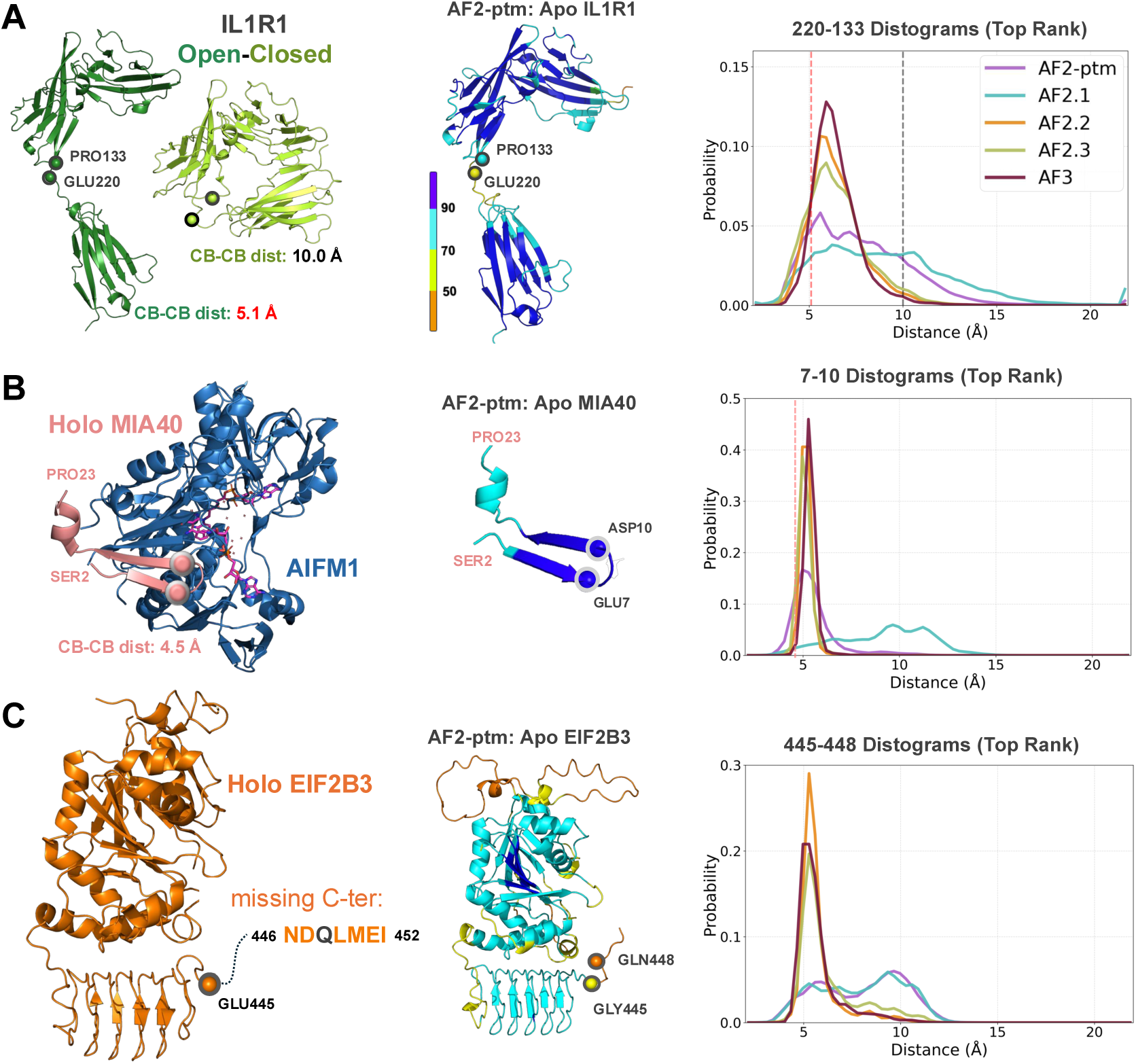
Distogram-based assessment of conformational flexibility across distinct structural contexts: **(A) IL1R1:** Comparison of existing IL1R1 capturing the open-to-closed transition. The left panel shows the corresponding structures for the open (dark green, PDB ID:1IRA_Y) and closed (light green, PDB ID:1G0Y_A) conformations, with the measured inter-residue distances (5.1 Å and 10.0 Å). **(B) MIA40:** The experimental structure of MIA40 in complex with AIFM1, sharing a similar beta-sheet stacking mode to that seen in the AK2/AIFM1 complex (PDB ID: 9GQZ). **(C) EIF2B3:** The holo EIF2B3 cryo-EM structure (PDB ID: 9HVD_A), highlighting the unresolved C-terminal sequence (residues 446–452). The central panel shows the AF2-ptm predicted apo structures colored by pLDDT, where blue indicates high confidence and orange indicates low confidence. For all cases, studied residues are emphasized with black or red spheres on the structures. The right panel depicts the distograms for the interested residue pairs across AlphaFold versions (best rankings only), with vertical dashed lines indicating the experimental reference distances if available.

The Interleukin IL1R1 receptor is a well-studied protein known to undergo a large-scale domain motion upon binding (PDB IDs: 1IRA, 1G0Y) [54,55]. In this case, we selected the 133-220 residue pair as our marker distance, which varies from 5.1 Å (open) to 10.0 Å (closed) (Figure 4A, S6A). Among all AF distograms generated, AF2-ptm and AF2.1 provided signals corresponding to both open and closed marker distances. Though, these profiles did not exhibit a clear bimodal pattern; instead, they displayed broad distributions skewed toward the closed-state distance. The other AF versions produced a sharp peak centered on the open state. Interestingly, each AF version predicted a different degree of domain closure at the structural level, even though both IL1R1 states were present during AF training (Figure S6D). When we ran MinnieFold in the template-free mode, the resulting distogram profiles largely reflected the same behavior, except for a few broad AF2.2 and AF2.3 distributions (Figure S7A). MSA clustering led to the peak shifts towards larger distances, accompanied by an increased probability in the final distogram bin (>22 Å). Relatedly, in the AF-derived structural ensembles, AF_cluster and AF3_cluster sampled both the open and closed marker distances, though AF_cluster also produced unrealistically long distances (Figure S7B). Among the generative models, Boltz-2 sampled both states, while BioEmu captured the closed state and Chai-1 sampled the open state (Figure S7B).

Our second benchmark protein, the oxidoreductase MIA40, binds to AIFM1 through its unstructured N-terminal region (resolved by NMR, PDB ID: 2K3J) [56]. Upon binding to AIFM1, this region adopts a structured conformation and forms a β-stacking interaction similar to that observed in AK2 (PDB ID: 9GQZ, Figure 4B) [41]. Surprisingly, all AF versions except AF2.1 confidently predicted this N-terminal region in its bound conformation, despite the fact that the holo MIA40 complex was not included in the AF training set (Figure S6D). To monitor binding-induced changes in the N-terminus, we used the distance between residues 7 and 10 as a marker (4.5 Å in the bound state). As anticipated, only the AF2.1 distogram displayed a broad marker distance distribution (Figure 4B, right). AF2-ptm exhibited a right-tailed distribution, which was amplified in MinnieFold runs, resulting in a secondary peak around 10 Å (Figure S8A). In this case, MSA column masking produced a mix of unimodal and bimodal peaks, whereas MSA clustering yielded multiple peaks. Within the AF-based ensembles, only AFsample2, AF_cluster, and AF3_cluster sampled both distance states, although the bound state remained dominant relative to the 10 Å state (Figure S8B). This behavior was similarly captured by BioEmu and Boltz-2.

Finally, we examined the eukaryotic translation initiation factor EIF2B3, which contains a C-terminal region (residues 446–452) that is unresolved in available cryo-EM structures (Figure 4C) [57]. All-to-all distograms of this region exhibited bimodality, where we selected the 445 and 448 inter-residue distance as our marker (Figure S6C). Since no structural information is available for this C-terminal tail, we focused primarily on the marker distance peak profiles. To this end, AF2.1 and AF2-ptm produced bimodal distributions, whereas other versions displayed a single peak centered at 5 Å, accompanied by a low-probability tail extending toward higher distances (Figure 4C). Under MinnieFold sampling, AF2.2 and AF2.3 showed a slight increase in probability within this long-distance tail (Figure S9A). Consistent with the other systems, MSA perturbation methods altered the distogram distributions, yielding either bimodal profiles or unimodal peaks (Figure S9A). Most ensemble-generation approaches successfully sampled bimodal distances, with the exceptions of AF3-multiseeds and AF3sample2 (Figure S9B). Collectively, these structural insights and the shared sequence motif with AK2’s C-tail (DQLMEI/DLVMFI) suggest a conserved, binding-induced hinge motion also for the EIF2B3 tail.

## Discussion

In this Perspective piece, we investigated whether AF-generated distograms can capture functionally relevant binding-induced hinge motions. For this, we focused on the AIFM1-induced structural change in AK2. Our findings demonstrate that AF2.3 and AF3 distograms for apo AK2 accurately recapitulate AK2’s binding-induced hinge motion, aligning closely with the MD-derived distance distributions. To evaluate broader applicability, we extended this analysis to three additional protein systems. These cases further supported the potential of AF-generated distograms to detect binding-induced hinge motions. Through the analysis of these cases, we also showed that not only bimodality, but also peak broadness may indicate hinge motion. Interestingly, earlier AF versions (AF2-ptm and AF2.1) appear more sensitive to initial flexibility detection. At the structural level, for all cases, the most realistic population profiles were observed in the AF3_cluster and Boltz2 ensembles.

These results suggest that for AF to model alternative distances, residue pairs must coevolve under competing evolutionary pressures, encoding structural heterogeneity. In the cases analyzed here, standard MMseqs2-generated MSAs were sufficient to track the expected structural changes. However, identifying more subtle or transient conformational states in other proteins may require further optimization of the input MSA. This is particularly critical for MSA clustering methods, where sequence depth significantly influences the distograms derived from reduced profiles. Indeed, in our dataset, IL1R1 (1,919 sequences) and MIA40 (3,487 sequences) possessed notably shallower MSAs compared to AK2 (17,000) and EIF2B3 (9,080), a disparity that likely contributed to the generation of unrealistic long-range distances in the former. In accordance with this, Audagnotto *et al.* also demonstrated that MSA generation strategies can significantly impact distogram quality [56]. In their work, they compared different MSA construction methods and assessed the resulting trRosetta distograms. Their results showed that DeepMSA produced more accurate distance distributions for myoglobin, effectively capturing both open and closed state conformations. However, these observations should be revisited by using AF, as distogram generation was not accessible at the time of their study.

As another MSA manipulation approach, AlphaFlex combines column masking with predicted residue flexibility, further illustrating how MSAs can be systematically optimized to sample conformational diversity [59]. AlphaFlex estimates residue-level flexibility by integrating structural, evolutionary, and physicochemical features. These flexibility profiles are then used to guide targeted MSA masking, selectively weakening coevolutionary constraints to generate physically plausible conformational states. While the concept is promising, it could not yet be tested for our test systems, as the implementation code is to be unpublished. Finally, probabilistic sampling methods such as bAIes and AlphaFold-Metainference [60–62] also utilize predicted distance distributions as restraints and have shown utility in modeling intrinsically disordered proteins (IDPs), highlighting the growing interest in distogram-based flexibility modeling.

Looking ahead, we suggest that AF distograms can be integrated into cryo-EM workflows as a pre-modeling diagnostic to mark flexible regions before atomic model building. This could help prioritize dynamic domains and guide targeted refinement efforts. A promising direction for future development would involve training strategies that automatically distinguish binding-induced changes from apo conformations. For this, one potential approach would be to identify specific MSA positions responsible for encoding binding-associated flexibility signals in distograms, informed by a systematic analysis of proteins with known bound–unbound landscapes. Though, to realize this potential, AF’s distogram resolution may need refinement. Current limitations, such as the fixed bin size of 0.375 Å and a maximum distance cap around 22 Å, can obscure large-scale transitions. Adaptive binning or fine-tuning strategies could help overcome these constraints and improve the utility of distograms for detecting conformational diversity relevant to cryo-EM interpretation.

## Conclusion

Overall, our findings show that while AF-derived confidence metrics, i.e., as pLDDT, PAE, and pTM, have been widely used to estimate protein flexibility [9,10, 63–67], distograms, which are generated prior to structure prediction, may encode a more fundamental layer of conformational information. Although preliminary, our results suggest that distogram profiles can act as effective, structure-free indicators of binding-induced hinge motions. As such, they hold promise for supporting the interpretation of flexible or partially resolved regions in cryo-EM maps. To enable follow-up studies and reproducibility, all data and scripts used in this work are publicly available at: https://github.com/CSB-KaracaLab/af-distogram-flexibility

## Author Contributions

BS: Conceptualization; Investigation; Writing - original draft; Writing - review & editing Methodology; Visualization; Formal analysis.

ABB: Investigation; Formal analysis; Methodology; Writing - original draft.

EK: Writing - original draft; Writing - review & editing; Project administration; Supervision; Conceptualization; Investigation; Methodology; Funding acquisition.

## Acknowledgments

This work was supported by the European Molecular Biology Organization (EMBO, grant IG4421) and the Scientific and Technological Research Council of Türkiye (TÜBİTAK, grant 122N790). We also acknowledge the Barcelona Supercomputing Center (BSC) and TÜBİTAK for providing computational resources on the MareNostrum5 supercomputing cluster, which were utilized for the AlphaFold3 runs. Access to MareNostrum5 was provided through a national access call coordinated by TÜBİTAK. We thank the reviewers for their constructive comments and valuable suggestions that helped improve this work.

## Abbreviation List

AF: AlphaFold
AIFM1: Apoptosis-Inducing Factor Mitochondrial 1
AK2: Adenylate Kinase 2
α-helix: Alpha helix
β: Beta
CAPRI: Critical Assessment of Predicted Interactions
Cβ: Beta carbon atom
cryo-EM: cryo-electron microscopy
DMPFold: DeepMetaPSICOV FolD
EIF2B3: Eukaryotic Translation Initiation Factor 2B Subunit 3
IDP: Intrinsically Disordered Protein
IL1R1: Interleukin-1 Receptor Type 1
MD: Molecular Dynamics
MIA40: Mitochondrial Intermembrane Space Import and Assembly Protein 40
MSA: Multiple Sequence Alignment
PAE: Predicted Aligned Error
PDB: Protein Data Bank
pLDDT: Predicted Local Distance Difference Test
pTM: Predicted TM-Score
RMSD: Root Mean Square Deviation
SI: Supplementary Information
Å: Ångström

## REFERENCES

1 . Karaca, E., & Bonvin, A. M. (2013). Advances in integrative modeling of biomolecular complexes. Methods (San Diego, Calif.), 59(3), 372–381. 10.1016/j.ymeth.2012.12.004

2. Bullock, J. M. A., Sen, N., Thalassinos, K., & Topf, M. (2018). Modeling Protein Complexes Using Restraints from Crosslinking Mass Spectrometry. Structure (London, England : 1993), 26(7), 1015–1024.e2. 10.1016/j.str.2018.04.016

3. Graziadei, A., & Rappsilber, J. (2022). Leveraging crosslinking mass spectrometry in structural and cell biology. Structure (London, England : 1993), 30(1), 37–54. 10.1016/j.str.2021.11.007

4 . Karaca, E., Rodrigues, J. P. G. L. M., Graziadei, A., Bonvin, A. M. J. J., & Carlomagno, T. (2017). M3: an integrative framework for structure determination of molecular machines. Nature methods, 14(9), 897–902. 10.1038/nmeth.4392 5

5. Skalidis, I., Kyrilis, F. L., Tüting, C., Hamdi, F., Chojnowski, G., & Kastritis, P. L. (2022). Cryo-EM and artificial intelligence visualize endogenous protein community members. Structure (London, England : 1993), 30(4), 575–589.e6. 10.1016/j.str.2022.01.001

6. Träger, T. K., Tüting, C., & Kastritis, P. L. (2024). The human touch: Utilizing AlphaFold 3 to analyze structures of endogenous metabolons. Structure (London, England : 1993), 32(10), 1555–1562. 10.1016/j.str.2024.08.018

7 . Jumper, J., Evans, R., Pritzel, A., Green, T., Figurnov, M., Ronneberger, O., Tunyasuvunakool, K., Bates, R., Žídek, A., Potapenko, A., Bridgland, A., Meyer, C., Kohl, S. A. A., Ballard, A. J., Cowie, A., Romera-Paredes, B., Nikolov, S., Jain, R., Adler, J., Back, T., … Hassabis, D. (2021). Highly accurate protein structure prediction with AlphaFold. Nature, 596(7873), 583–589. 10.1038/s41586-021-03819-2

8 . Mosalaganti, S., Obarska-Kosinska, A., Siggel, M., Taniguchi, R., Turoňová, B., Zimmerli, C. E., Buczak, K., Schmidt, F. H., Margiotta, E., Mackmull, M. T., Hagen, W. J. H., Hummer, G., Kosinski, J., & Beck, M. (2022). AI-based structure prediction empowers integrative structural analysis of human nuclear pores. Science (New York, N.Y.), 376(6598), eabm9506. 10.1126/science.abm9506

9 . Saldaño, T., Escobedo, N., Marchetti, J., Zea, D. J., Mac Donagh, J., Velez Rueda, A. J., Gonik, E., García Melani, A., Novomisky Nechcoff, J., Salas, M. N., Peters, T., Demitroff, N., Fernandez Alberti, S., Palopoli, N., Fornasari, M. S., & Parisi, G. (2022). Impact of protein conformational diversity on AlphaFold predictions. *Bioinformatics (Oxford*, England), 38(10), 2742–2748. 10.1093/bioinformatics/btac202

10 . Carugo, O. (2023). pLDDT Values in AlphaFold2 Protein Models Are Unrelated to Globular Protein Local Flexibility. Crystals, 13(11), 1560. 10.3390/cryst13111560

11 . Del Alamo, D., Sala, D., Mchaourab, H. S., & Meiler, J. (2022). Sampling alternative conformational states of transporters and receptors with AlphaFold2. eLife, 11, e75751. 10.7554/eLife.75751

12 . Guo, H. B., Perminov, A., Bekele, S., Kedziora, G., Farajollahi, S., Varaljay, V., Hinkle, K., Molinero, V., Meister, K., Hung, C., Dennis, P., Kelley-Loughnane, N., & Berry, R. (2022). AlphaFold2 models indicate that protein sequence determines both structure and dynamics. Scientific reports, 12(1), 10696. 10.1038/s41598-022-14382-9

13 . Tüting, C., Schmidt, L., Skalidis, I., Sinz, A., & Kastritis, P. L. (2023). Enabling cryo-EM density interpretation from yeast native cell extracts by proteomics data and AlphaFold structures. Proteomics, 23(17), e2200096. 10.1002/pmic.202200096

14 . Jiang, T., Thielges, M. C., & Feng, C. (2025). Emerging approaches to investigating functional protein dynamics in modular redox enzymes: Nitric oxide synthase as a model system. The Journal of biological chemistry, 301(3), 108282. 10.1016/j.jbc.2025.108282

15 . He, Y., Haque, M. M., Stuehr, D. J., & Lu, H. P. (2021). Conformational states and fluctuations in endothelial nitric oxide synthase under calmodulin regulation. Biophysical journal, 120(23), 5196–5206. 10.1016/j.bpj.2021.11.001

16 . Tüting, C., Schmidt, L., Skalidis, I., Sinz, A., & Kastritis, P. L. (2023). Enabling cryo-EM density interpretation from yeast native cell extracts by proteomics data and AlphaFold structures. Proteomics, 23(17), e2200096. 10.1002/pmic.202200096

17 . Skalidis, I., Kyrilis, F. L., Tüting, C., Hamdi, F., Träger, T. K., Belapure, J., Hause, G., Fratini, M., O’Reilly, F. J., Heilmann, I., Rappsilber, J., & Kastritis, P. L. (2023). Structural analysis of an endogenous 4-megadalton succinyl-CoA-generating metabolon. Communications biology, 6(1), 552. 10.1038/s42003-023-04885-0

18. Skalidis, I., Kyrilis, F. L., Tüting, C., Hamdi, F., Chojnowski, G., & Kastritis, P. L. (2022). Cryo-EM and artificial intelligence visualize endogenous protein community members. Structure (London, England : 1993), 30(4), 575–589.e6. 10.1016/j.str.2022.01.001

19 . Kyrilis, F. L., Semchonok, D. A., Skalidis, I., Tüting, C., Hamdi, F., O’Reilly, F. J., Rappsilber, J., & Kastritis, P. L. (2021). Integrative structure of a 10-megadalton eukaryotic pyruvate dehydrogenase complex from native cell extracts. Cell reports, 34(6), 108727. 10.1016/j.celrep.2021.108727

20 . Forsberg, B. O., Aibara, S., Howard, R. J., Mortezaei, N., & Lindahl, E. (2020). Arrangement and symmetry of the fungal E3BP-containing core of the pyruvate dehydrogenase complex. Nature communications, 11(1), 4667. 10.1038/s41467-020-18401-z

21 . Tüting, C., Kyrilis, F. L., Müller, J., Sorokina, M., Skalidis, I., Hamdi, F., Sadian, Y., & Kastritis, P. L. (2021). Cryo-EM snapshots of a native lysate provide structural insights into a metabolon-embedded transacetylase reaction. Nature communications, 12(1), 6933. 10.1038/s41467-021-27287-4

22 . Cui, X., Ge, L., Chen, X., Lv, Z., Wang, S., Zhou, X., & Zhang, G. (2025). Beyond static structures: protein dynamic conformations modeling in the post-AlphaFold era. Briefings in bioinformatics, 26(4), bbaf340. 10.1093/bib/bbaf340

23 . Stein, R. A., & Mchaourab, H. S. (2022). SPEACH_AF: Sampling protein ensembles and conformational heterogeneity with Alphafold2. PLoS computational biology, 18(8), e1010483. 10.1371/journal.pcbi.1010483

24 . Wallner B. (2023). AFsample: improving multimer prediction with AlphaFold using massive sampling. Bioinformatics (Oxford, England), 39(9), btad573. 10.1093/bioinformatics/btad573

25 . Raouraoua, N., Mirabello, C., Véry, T., Blanchet, C., Wallner, B., Lensink, M. F., & Brysbaert, G. (2024). MassiveFold: unveiling AlphaFold’s hidden potential with optimized and parallelized massive sampling. Nature computational science, 4(11), 824–828. 10.1038/s43588-024-00714-4

26 . Jing B, Berger B, Jaakkola A. (2024) AlphaFold Meets Flow Matching for Generating Protein Ensembles. Rxiv. 10.48550/arXiv.2402.04845

27 . Kalakoti, Y., & Wallner, B. (2025). AFsample2 predicts multiple conformations and ensembles with AlphaFold2. Communications biology, 8(1), 373. 10.1038/s42003-025-07791-9

28 . Wayment-Steele, H. K., Ojoawo, A., Otten, R., Apitz, J. M., Pitsawong, W., Hömberger, M., Ovchinnikov, S., Colwell, L., & Kern, D. (2024). Predicting multiple conformations via sequence clustering and AlphaFold2. Nature, 625(7996), 832–839. 10.1038/s41586-023-06832-9

29 . Sala, D., Hildebrand, P. W., & Meiler, J. (2023). Biasing AlphaFold2 to predict GPCRs and kinases with user-defined functional or structural properties. Frontiers in molecular biosciences, 10, 1121962. 10.3389/fmolb.2023.1121962

30 . Heo, L., & Feig, M. (2022). Multi-state modeling of G-protein coupled receptors at experimental accuracy. Proteins, 90(11), 1873–1885. 10.1002/prot.26382

31 . Schwarz, D., Georges, G., Kelm, S., Shi, J., Vangone, A., & Deane, C. M. (2021). Co-evolutionary distance predictions contain flexibility information. Bioinformatics (Oxford, England), 38(1), 65–72. 10.1093/bioinformatics/btab562

32 . Senior, A. W., Evans, R., Jumper, J., Kirkpatrick, J., Sifre, L., Green, T., Qin, C., Žídek, A., Nelson, A. W. R., Bridgland, A., Penedones, H., Petersen, S., Simonyan, K., Crossan, S., Kohli, P., Jones, D. T., Silver, D., Kavukcuoglu, K., & Hassabis, D. (2020). Improved protein structure prediction using potentials from deep learning. Nature, 577(7792), 706–710. 10.1038/s41586-019-1923-7

33 . Greener, J. G., Kandathil, S. M., & Jones, D. T. (2019). Deep learning extends de novo protein modelling coverage of genomes using iteratively predicted structural constraints. Nature communications, 10(1), 3977. 10.1038/s41467-019-11994-0

34 . Schildhauer, F., Ryl, P. S. J., Lauer, S. M., Lenz, S., Barlas, A. B., Ouzounidis, V. R., Jeffrey, K., Marcu, D. C., O’Reilly, F. J., Graziadei, A., Stuiver, M., Schmidt, K., Ewers, H., Spahn, C. M. T., Karaca, E., Busch, K. E., Cheerambathur, D., Schwefel, D., & Rappsilber, J. (2025). An NADH-controlled gatekeeper of ATP synthase. Molecular cell, 85(13), 2567–2580.e12. 10.1016/j.molcel.2025.06.007

35 . Abramson, J., Adler, J., Dunger, J., Evans, R., Green, T., Pritzel, A., Ronneberger, O., Willmore, L., Ballard, A. J., Bambrick, J., Bodenstein, S. W., Evans, D. A., Hung, C. C., O’Neill, M., Reiman, D., Tunyasuvunakool, K., Wu, Z., Žemgulytė, A., Arvaniti, E., Beattie, C., … Jumper, J. M. (2024). Accurate structure prediction of biomolecular interactions with AlphaFold 3. Nature, 630(8016), 493–500. 10.1038/s41586-024-07487-w

36 . Burkart, A., Shi, X., Chouinard, M., & Corvera, S. (2011). Adenylate kinase 2 links mitochondrial energy metabolism to the induction of the unfolded protein response. The Journal of biological chemistry, 286(6), 4081–4089. 10.1074/jbc.M110.134106

37 . Dzeja, P., & Terzic, A. (2009). Adenylate Kinase and AMP Signaling Networks: Metabolic Monitoring, Signal Communication and Body Energy Sensing. International Journal of Molecular Sciences, 10(4), 1729–1772. 10.3390/ijms10041729

38 . Formoso, E., Limongelli, V., & Parrinello, M. (2015). Energetics and structural characterization of the large-scale functional motion of adenylate kinase. Scientific reports, 5, 8425. 10.1038/srep08425

39 . Snow, C., Qi, G., & Hayward, S. (2007). Essential dynamics sampling study of adenylate kinase: comparison to citrate synthase and implication for the hinge and shear mechanisms of domain motions. Proteins, 67(2), 325–337. 10.1002/prot.21280

40 . Whitford, P. C., Gosavi, S., & Onuchic, J. N. (2008). Conformational transitions in adenylate kinase. Allosteric communication reduces misligation. The Journal of biological chemistry, 283(4), 2042–2048. 10.1074/jbc.M707632200

41 . Rothemann, R. A., Pavlenko, E., Mondal, M., Gerlich, S., Grobushkin, P., Mostert, S., Racho, J., Weiss, K., Stobbe, D., Stillger, K., Lapacz, K., Salscheider, S. L., Petrungaro, C., Ehninger, D., Nguyen, T. H. D., Dengjel, J., Neundorf, I., Bano, D., Poepsel, S., & Riemer, J. (2025). Interaction with AK2A links AIFM1 to cellular energy metabolism. Molecular cell, 85(13), 2550–2566.e6. 10.1016/j.molcel.2025.05.036

42. 2C9Y X-ray structure, (2005). 10.2210/pdb2C9Y/pdb

43. Barlas, A.B. (2024). Understanding the function of biomolecular complexes through the lens of molecular dynamics. [Doctoral dissertation, Dokuz Eylul University, Izmir]. Source: https://tez.yok.gov.tr/UlusalTezMerkezi/TezGoster?key=E_eEUHQic_C-LvhxNQn1W3wmOCInlBnVAZxdJWqWdN5YkWRUeQ_5_WUDFHGmyRtK

44 . Steinegger, M., & Söding, J. (2017). MMseqs2 enables sensitive protein sequence searching for the analysis of massive data sets. Nature biotechnology, 35(11), 1026–1028. 10.1038/nbt.3988

45 . Mirdita, M., Schütze, K., Moriwaki, Y., Heo, L., Ovchinnikov, S., & Steinegger, M. (2022). ColabFold: making protein folding accessible to all. Nature methods, 19(6), 679–682. 10.1038/s41592-022-01488-1

46 . Savaş, B., Yılmazbilek, İ., Özsan, A., & Karaca, E. (2025). Towards a Greener AlphaFold2 Protocol for Antibody-Antigen Modeling: Insights From CAPRI Round 55. Proteins, 10.1002/prot.26820. Advance online publication. https://doi.org/10.1002/prot.26820

47 . Evans R, O’Neill M, Pritzel A, Antropova N, Senior A, Green T, Žídek A, Bates R, Blackwell S, Yim J, et al. (2022) Protein complex prediction with AlphaFold-multimer. bioRxiv. 10.1101/2021.10.04.463034

48. https://github.com/google-deepmind/alphafold/blob/main/docs/technical_note_v2.3.0.md AlphaFold v2.3 technical note, released after CASP15.

49 . Ozden, B., Kryshtafovych, A., & Karaca, E. (2023). The impact of AI-based modeling on the accuracy of protein assembly prediction: Insights from CASP15. Proteins, 91(12), 1636–1657. 10.1002/prot.26598

50 . Guan, L., & Keating, A. E. (2025). Training bias and sequence alignments shape protein-peptide docking by AlphaFold and related methods. Protein science : a publication of the Protein Society, 34(11), e70331. 10.1002/pro.70331

51 . Lewis, S., Hempel, T., Jiménez-Luna, J., Gastegger, M., Xie, Y., Foong, A. Y. K., Satorras, V. G., Abdin, O., Veeling, B. S., Zaporozhets, I., Chen, Y., Yang, S., Foster, A. E., Schneuing, A., Nigam, J., Barbero, F., Stimper, V., Campbell, A., Yim, J., Lienen, M., … Noé, F. (2025). Scalable emulation of protein equilibrium ensembles with generative deep learning. Science (New York, N.Y.), 389(6761), eadv9817. 10.1126/science.adv9817

52 . Chai Discovery, Boitreaud, J., Dent, J., McPartlon, M., Meier, J., Reis, V., Rogozhnikov, A., & Wu, K. (2024). Chai-1: Decoding the molecular interactions of life. bioRxiv. 10.1101/2024.10.10.615955

53 . Passaro, S., Corso, G., Wohlwend, J., Reveiz, M., Thaler, S., Somnath, V. R., Getz, N., Portnoi, T., Roy, J., Stark, H., Kwabi-Addo, D., Beaini, D., Jaakkola, T., & Barzilay, R. (2025). Boltz-2: Towards Accurate and Efficient Binding Affinity Prediction. bioRxiv. 10.1101/2025.06.14.659707

54 . Schreuder, H., Tardif, C., Trump-Kallmeyer, S., Soffientini, A., Sarubbi, E., Akeson, A., Bowlin, T., Yanofsky, S., & Barrett, R. W. (1997). A new cytokine-receptor binding mode revealed by the crystal structure of the IL-1 receptor with an antagonist. Nature, 386(6621), 194–200. 10.1038/386194a0

55 . Vigers, G. P., Dripps, D. J., Edwards, C. K., 3rd, & Brandhuber, B. J. (2000). X-ray crystal structure of a small antagonist peptide bound to interleukin-1 receptor type 1. The Journal of biological chemistry, 275(47), 36927–36933. 10.1074/jbc.M006071200

56 . Banci, L., Bertini, I., Cefaro, C., Ciofi-Baffoni, S., Gallo, A., Martinelli, M., Sideris, D. P., Katrakili, N., & Tokatlidis, K. (2009). MIA40 is an oxidoreductase that catalyzes oxidative protein folding in mitochondria. Nature structural & molecular biology, 16(2), 198–206. 10.1038/nsmb.1553

57 . Circir, A., Koksal Bicakci, G., Savas, B., Doken, D. N., Henden, Ş. O., Can, T., Karaca, E., & Erson-Bensan, A. E. (2022). A C-term truncated EIF2Bγ protein encoded by an intronically polyadenylated isoform introduces unfavorable EIF2Bγ-EIF2γ interactions. Proteins, 90(3), 889–897. 10.1002/prot.26284

58. Audagnotto, M., Czechtizky, W., De Maria, L., Käck, H., Papoian, G., Tornberg, L., Tyrchan, C., & Ulander, J. (2022). Machine learning/molecular dynamic protein structure prediction approach to investigate the protein conformational ensemble. Scientific reports, 12(1), 10018. 10.1038/s41598-022-13714-z

59 . Ge, L., Cui, X., Zhao, K., Zhou, X., Zhang, Y., & Zhang, G. (2025). AlphaFlex: Accuracy modeling of protein multiple conformations via predicted flexible residues. bioRxiv. 10.1101/2025.07.11.664327

60 . Brotzakis, Z. F., Zhang, S., Murtada, M. H., & Vendruscolo, M. (2025). AlphaFold prediction of structural ensembles of disordered proteins. Nature communications, 16(1), 1632. 10.1038/s41467-025-56572-9

61 . Schnapka, V., Morozova, T. I., Sen, S., & Bonomi, M. (2025). Atomic resolution ensembles of intrinsically disordered proteins with Alphafold. bioRxiv. 10.1101/2025.06.18.660298

62 . Sen, S., Hoff, S. E., Morozova, T. I., Schnapka, V., & Bonomi, M. (2025). Advancing in silico drug design with Bayesian refinement of AlphaFold models. bioRxiv. 10.1101/2025.06.25.661454

63. Vander Meersche, Y., Diharce, J., Gelly, J. C., & Galochkina, T. (2025). Flexibility or uncertainty? A critical assessment of AlphaFold 2 pLDDT. Structure (London, England: 1993), S0969-2126(25)00344-2. Advance online publication. 10.1016/j.str.2025.09.001

64 . Ma, P., Li, D. W., & Brüschweiler, R. (2023). Predicting protein flexibility with AlphaFold. Proteins, 91(6), 847–855. 10.1002/prot.26471

65 . Wróblewski, K., & Kmiecik, S. (2024). Integrating AlphaFold pLDDT Scores into CABS-flex for enhanced protein flexibility simulations. Computational and structural biotechnology journal, 23, 4350–4356. 10.1016/j.csbj.2024.11.047

66 . Tesei, G., Trolle, A. I., Jonsson, N., Betz, J., Knudsen, F. E., Pesce, F., Johansson, K. E., & Lindorff-Larsen, K. (2024). Conformational ensembles of the human intrinsically disordered proteome. Nature, 626(8000), 897–904. 10.1038/s41586-023-07004-5

67 . Jussupow, A., & Kaila, V. R. I. (2023). Effective Molecular Dynamics from Neural Network-Based Structure Prediction Models. Journal of chemical theory and computation, 19(7), 1965–1975. 10.1021/acs.jctc.2c01027

