## Supporting Information for "Exploring the Potential of AlphaFold Distograms for Predicting Binding-induced Hinge Motions"

### **SUPPLEMENTARY TEXT**

1. **Data and Code Availability**

We share the multiple sequence alignments (MSAs), distogram outputs, predicted PDB files, code used to generate the figures in this manuscript, and additional useful scripts at <https://github.com/CSB-KaracaLab/af-distogram-flexibility>.

2. **Methods**

Pairwise distograms were extracted from the AF2 logits, converted into probability distributions using the softmax function across distance bins. The distograms consist of 64 bins ranging from 2.0 Å to >22.2 Å with a constant bin width of 0.3125 Å. Each bar in the predicted distograms and MD-derived distograms corresponds to these distance bins.

For MSA generation, we utilized MMseqs2 through local ColabFold, ensuring consistent alignment across all AlphaFold versions in unpaired MSA mode. In all modeling scenarios, the template usage is disabled to avoid influence of recently published AK2:AIFM1 structure.

AF2 is used through local ColabFold (version 1.5.5) on a lab-scale workstation equipped with an A4000 GPU. AF3 modelings were performed on the MareNostrum5 supercomputer.

For each figure, the methodology is detailed below.

**Figure 1**

Each version was run five times with different random seeds (starting from seed 0 and incrementing by 1). We used AF3 with default settings were applied for both versions.

**Figure 2 (A)**

We generated 25 distograms per AF version with random seeds (starting from seed 0 and incrementing by 1). Since AF3 generates one distogram per run, 25 separate runs were submitted. The alphafold_ptm weight unless stated differently.

**Figure 2 (B, C)**

MinnieFold run scenarios are summarized in Table S1, and the complete protocol is available at: <https://github.com/CSB-KaracaLab/MinnieFold>

**Table S1. MinnieFold scenario followed in this work.**

| **Scenario** | **AF2 Version** | **Dropout** | **Templates** | **Recycles** | **Models** |
| --- | --- | --- | --- | --- | --- |
| 1–4 | all | Yes | Yes | Default | 5x4 |
| 5–8 | all | Yes | No | Default | 5x4 |
| 9–12 | all | Yes | Yes | 9 | 5x4 |
| 13–16 | all | Yes | No | 9 | 5x4 |
| 17–20 | all | Yes | Yes | 21 | 5x4 |
| 21–24 | all | Yes | No | 21 | 5x4 |

AFsample2 was accessed from: <https://github.com/iamysk/AFsample2>. Following Kalakoti and Wallner (2025), we applied 15% random masking to MSAs. Distograms were generated using the alphafold_ptm weights over 500 masked MSAs. Three recycle steps were applied, and template usage and dropouts were disabled.

MSA clustering was performed using the ClusterMSA.py script from: <https://github.com/HWaymentSteele/AF_Cluster>. The resulting clustered MSAs were input into the local ColabFold pipeline with three recycle steps and no template usage. Modeling was performed using only the alphafold_ptm model1 weights, applied to a total of 407 clustered MSAs. Three recycle steps were applied, and template usage and dropouts were disabled.

**Figure 2 (D)**

We generated column masked MSAs from the “masked_msa” data stored in AFsample2 output, using our in-house column_masking.py script which masks the same amino acid positions masked during the AFsample2 generation.

For AF3_cluster, we used the same clustered MSAs as inputs for AF3 pipeline.

**Figure 3**

Since AF3 produces a single distogram and five structural models per run, we evaluated the first model (sample-1) from each run to ensure a consistent number of outputs between AF2 and AF3.

BioEmu was accessed through its standard interface using num_samples = 500, with filter_samples disabled; all other parameters were kept at their default settings. The corresponding Colab notebook is available at: <https://colab.research.google.com/github/sokrypton/ColabFold/blob/main/BioEmu.ipynb>

Boltz-2 was accessed via <https://colab.research.google.com/drive/1iXyJOz-kk9ncWyFDNC62HjSrVyk4yooi> [1]. The same MSA used throughout this study was applied. Default parameters were used, including diffusion_samples = 5, recycling_steps = 3, sampling_steps = 200, and max_msa_seqs = 8192. A total of 100 different random seeds were used, ranging from 0 to 99.

We ran Chai-1 using the same MSA as in the other analyses. The model was executed with num_trunk_recycles = 3, num_diffn_timesteps = 200, and seed numbers ranging from 0 to 99. The notebook is accessible at <https://colab.research.google.com/drive/1z3KOjbTHde2c4TaufMa2MGdeAnd5QPj_>.

3. **Supplementary Figures**

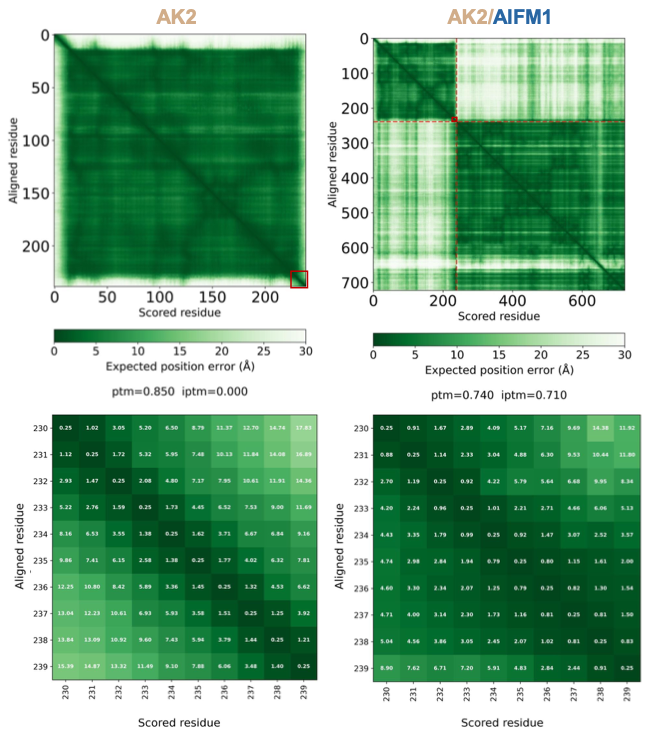

**Figure S1. Predicted aligned error (PAE) plots of AK2 (left) and the AK2/AIFM1 complex (right).** The color bar represents the expected position error (in Å) between residue pairs, with darker green indicating higher confidence and lighter regions denoting greater uncertainty. The bottom panels show zoomed-in PAE plots for the last 10 residues, with individual PAE scores displayed within each cell.

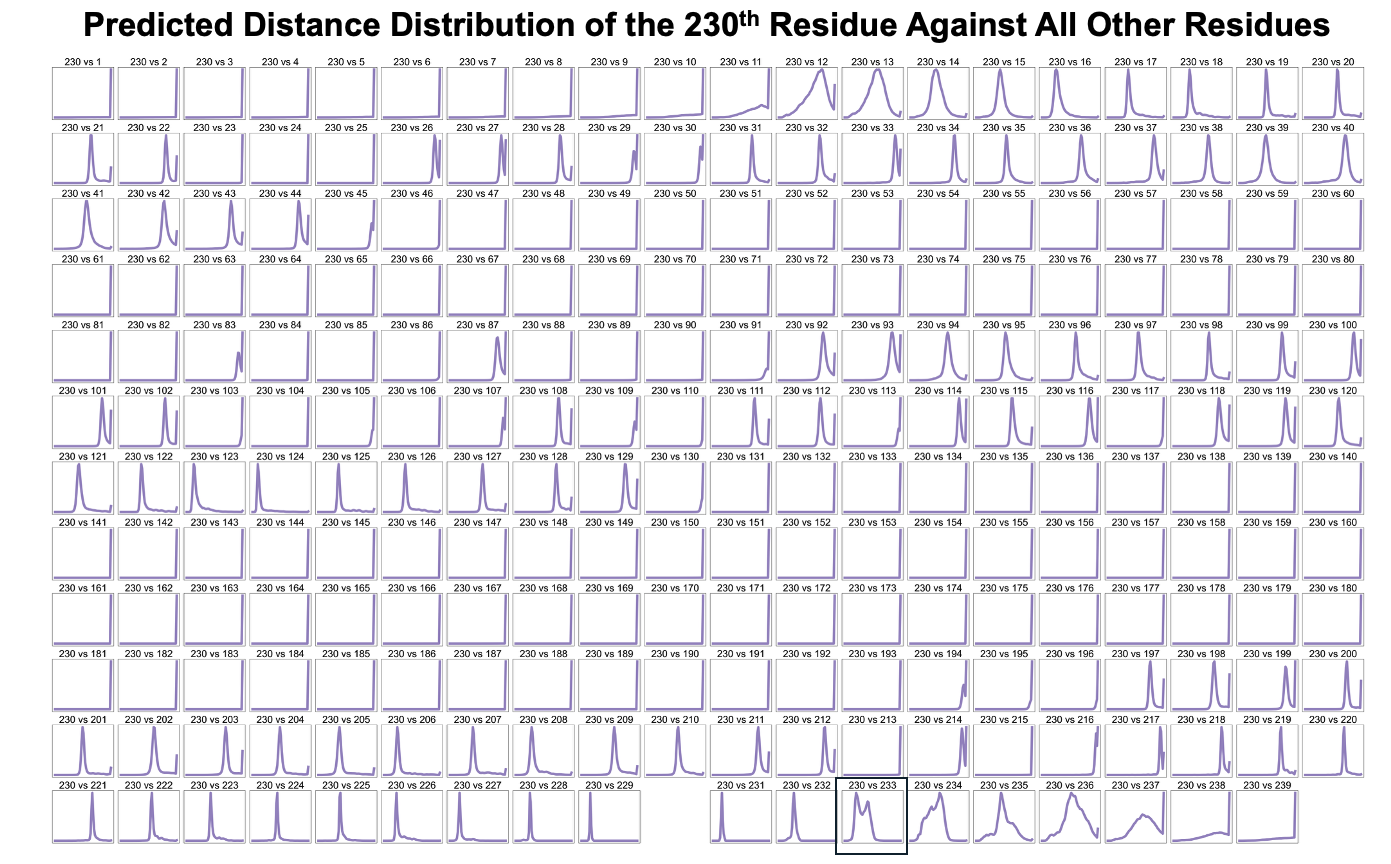

**Figure S2. Predicted distance probability profiles for residue 230 against all other residues in AK2.** Each subplot represents the AF2-ptm predicted distance distribution for residue 230 in pairwise comparison with every other residue in the sequence. For visual simplicity, the x-axis (distance) and y-axis (probability) are omitted. Note that the x-axis range may vary across subplots showing different distance distributions.

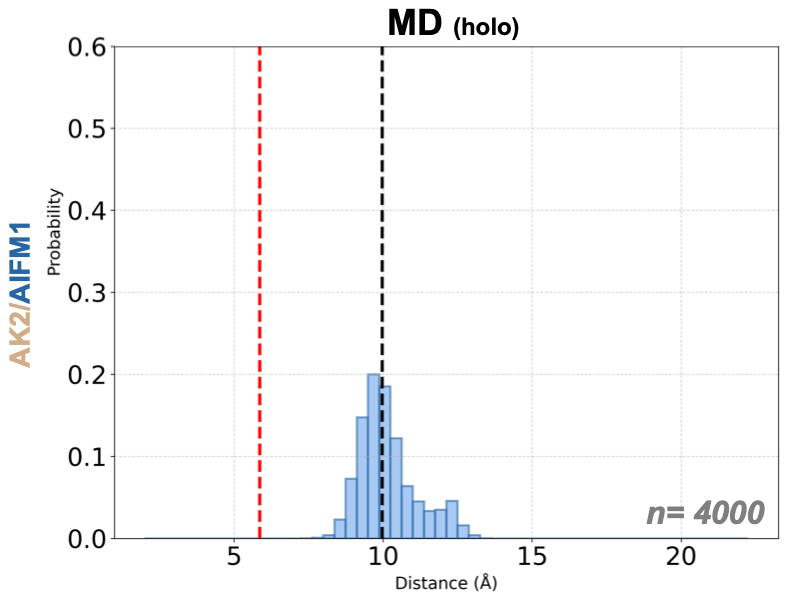

**Figure S3. The measured distance probability distribution, obtained from AK2/AIFM1 holo 4 μs all-atom MD simulations.** The dashed lines indicate the threshold distances obtained from MD simulations: at 6.0 Å in red for the apo and 10.0 Å in black for the holo state. The sample size for the plot is indicated with n.

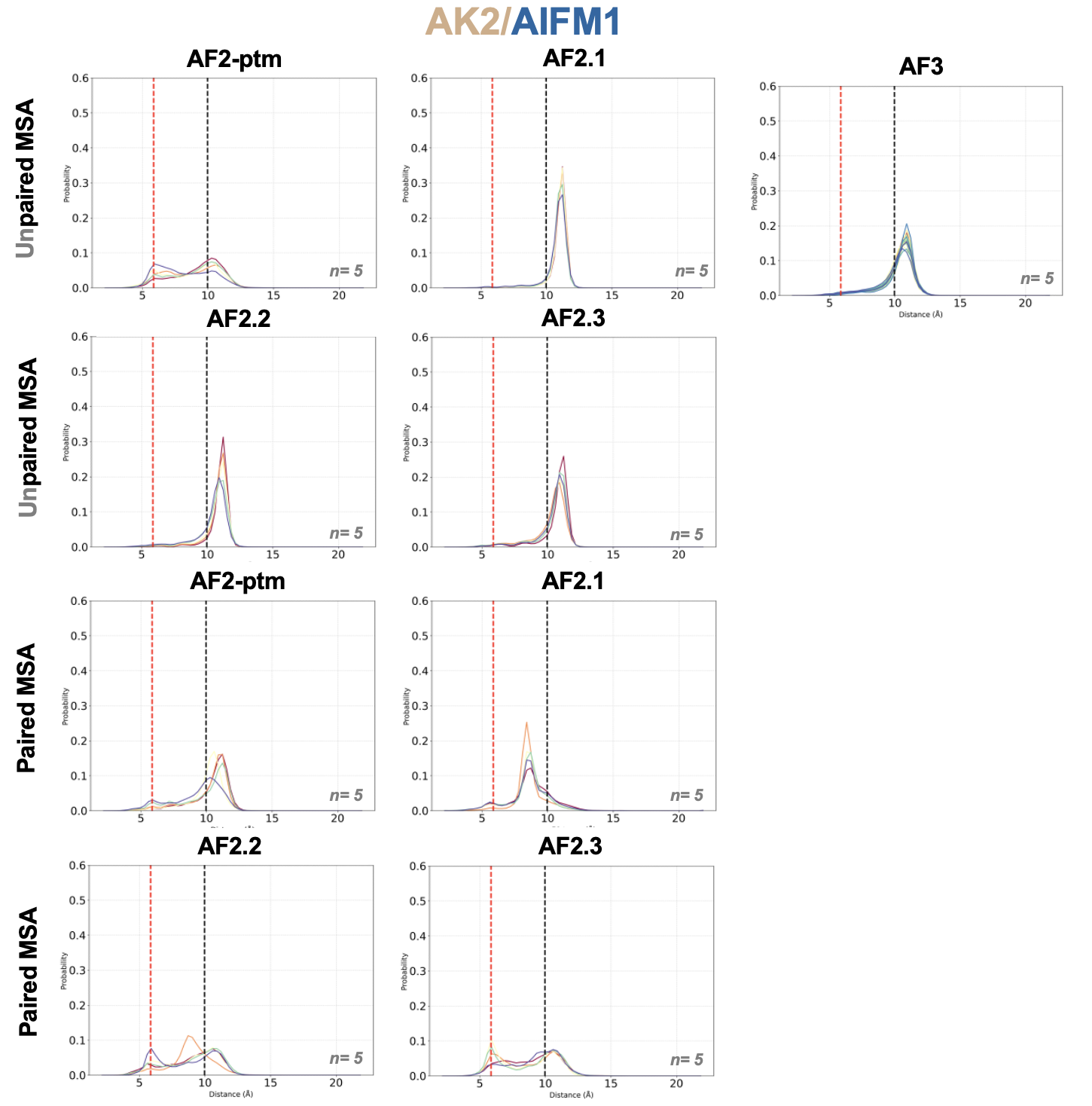

**Figure S4.** Distogram profiles of the predicted 230-233 marker distance of holo AK2/AIFM1. Distogram profiles of 230-233 pair calculated by all AlphaFold versions available using paired or unpaired MSAs. The dashed lines indicate the threshold distances obtained from MD simulations: at 6.0 Å in red for the apo and 10.0 Å in black for the holo state. The sample size for the plot is indicated with n. *AF3 pipeline could not be used with paired MSA.

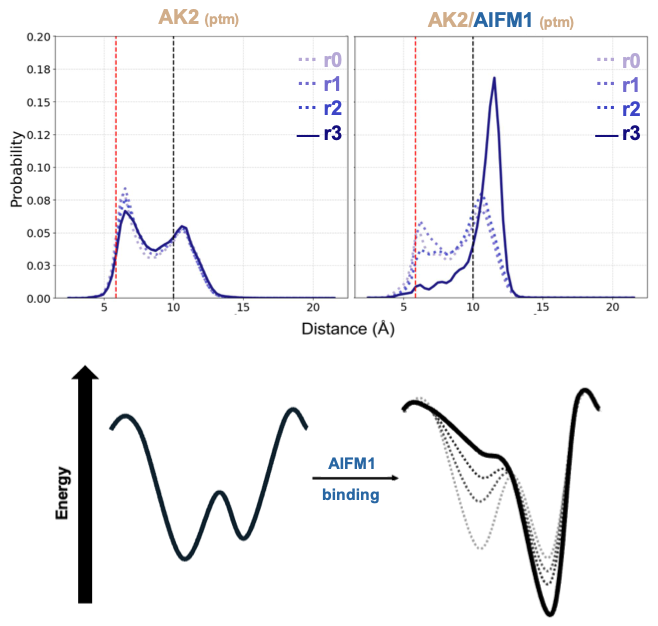

**Figure S5. Distogram profiles of the predicted 230-233 marker distance of apo and holo AK2 across three recycles**. Distogram profiles of 230-233 pair calculated by all AlphaFold2-ptm versions (paired and unpaired MSA, respectively). The dashed lines indicate the threshold distances obtained from MD simulations: at 6.0 Å in red for the apo and 10.0 Å in black for the holo state.

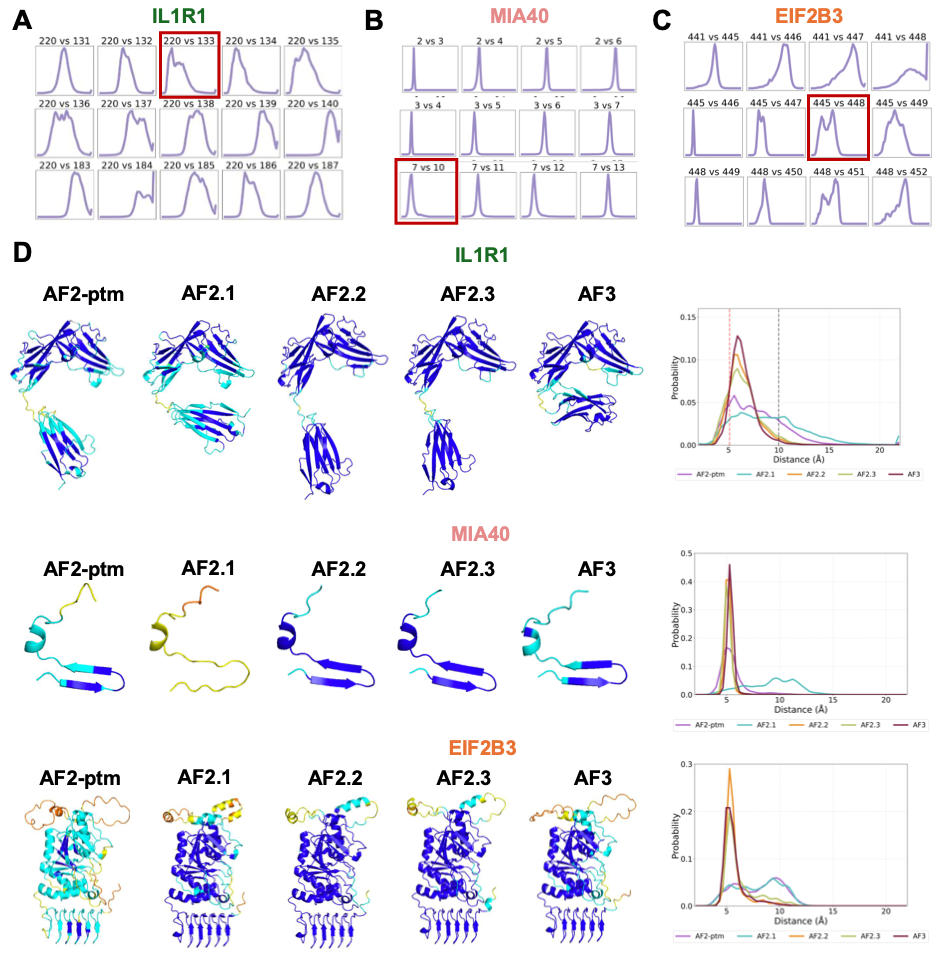

**Figure S6. All-to-all distogram analysis for IL1R1, MIA40 and EIF2B3 and distogram-pLDDT relation.** (A–C) Representative plots from all-to-all residue distograms. Selected pairwise interactions are displayed, with specific pairs of interest highlighted in red boxes. (D) The reflection of distogram profiles in output structures, evaluated with pLDDT scoring.

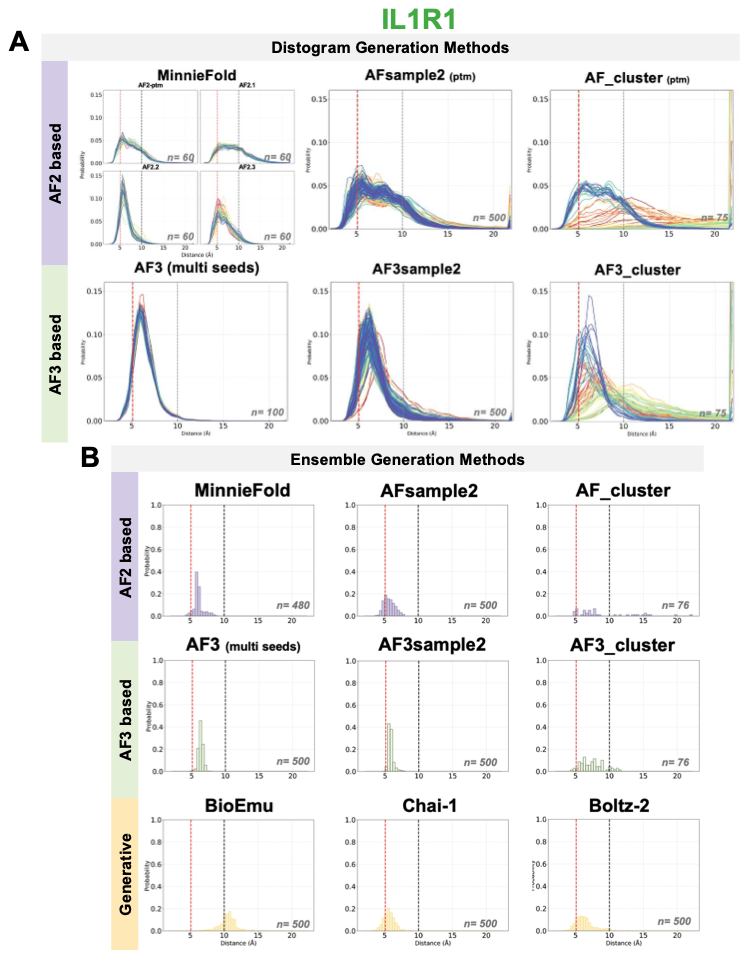

**Figure S7. AF-based distogram profiles (A) and measured distance distribution across ensemble generation tools (B) for IL1R1.** 133-220 residue pair is used for the analysis. The dashed lines indicate the distances obtained from open (5.1 Å, red) and closed (10.0 Å, black) states. The sample size for the plot is indicated with n.

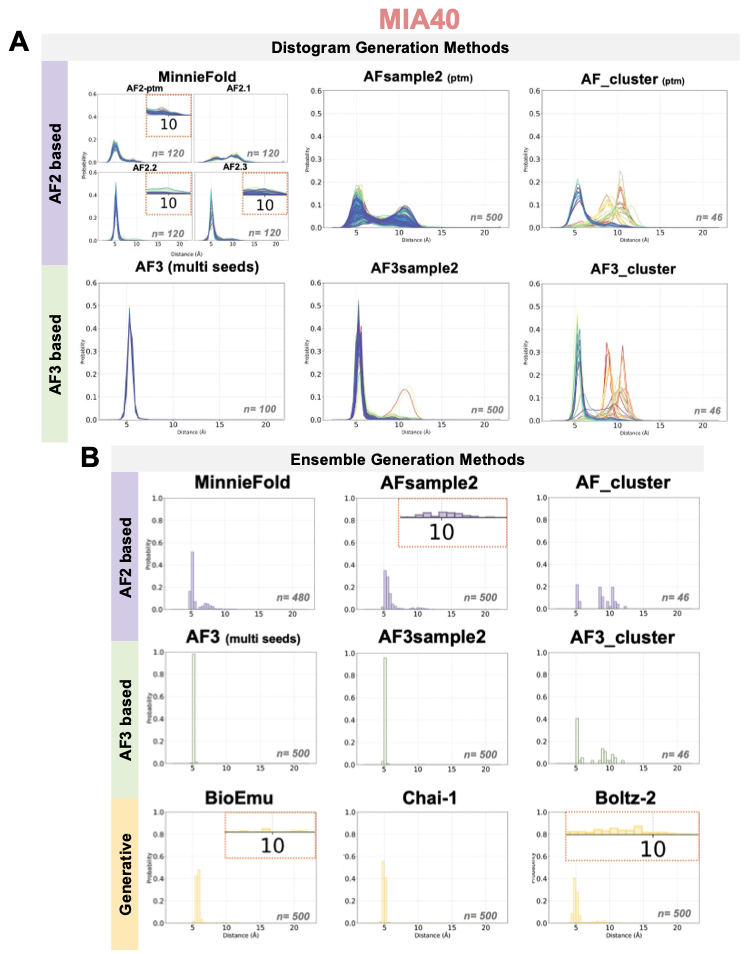
**Figure S8. AF-based distogram profiles (A) and measured distance distribution across ensemble generation tools (B) for MIA40.** 7-10 residue pair is used for the analysis. The sample size for the plot is indicated with n.

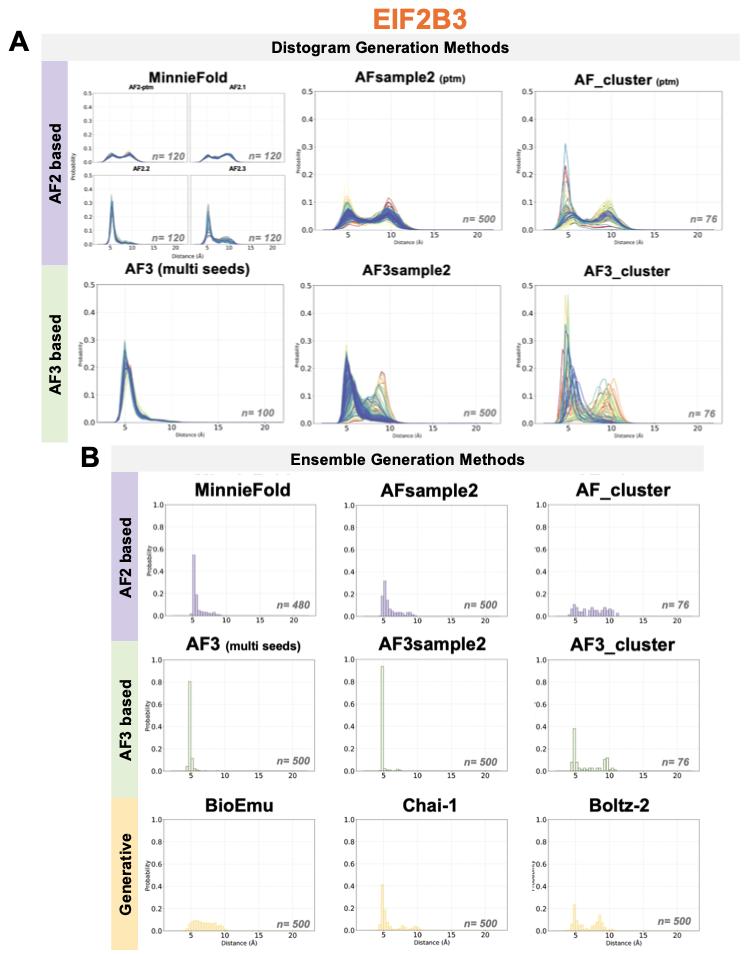

**Figure S9. AF-based distogram profiles (A) and measured distance distribution across ensemble generation tools (B) for EIF2B3.** 445-448 residue pair is used for the analysis. The sample size for the plot is indicated with n.

**SUPPLEMANTARY REFERENCES**

1. Kim, J.C. (2025). Computational Chemistry for Fun. Zenodo. <https://doi.org/10.5281/zenodo.14881401>.
